# Fast model-free standardization and integration of single-cell transcriptomics data

**DOI:** 10.1101/2022.03.28.486110

**Authors:** Yang Xu, Rafael Kramann, Rachel Patton McCord, Sikander Hayat

## Abstract

Single-cell transcriptomics datasets from the same anatomical sites generated by different research labs are becoming mainstream. However, fast, and computationally inexpensive tools for standardization of cell-type annotation and data integration are still needed to increase research inclusivity. To standardize cell-type annotation and integrate single-cell transcriptomics datasets, we have built a fast, model-free integration method called MASI (**M**arker-**A**ssisted **S**tandardization and **I**ntegration). MASI can run integrative annotation on a personal laptop for approximately one million cells, providing a cheap computational alternative for the single-cell data analysis community. MASI has an average macro F1/overall accuracy of 0.79/0.89 over the 4 benchmark datasets. We demonstrate that MASI outperforms other methods based on speed, and its performance for the tasks of data integration and cell-type annotation is comparable or even superior to other existing methods. We apply MASI for integrative lineage analysis and show that it preserves the underlying biological signal in datasets tested. Finally, to harness knowledge from single-cell atlases, we demonstrate three case studies that cover integration across research groups, biological conditions, and surveyed participants, respectively.

## Introduction

Single-cell RNA-seq (scRNA-seq) technologies have rapidly evolved over the last decade. Numerous studies have demonstrated the utility of single-cell transcriptomics datasets in improving our understanding of cellular heterogeneity and molecular mechanisms at unprecedented resolution. Over the past years, many single-cell datasets have been made available from different research groups, using multiple single-cell platforms, and covering diverse biological conditions. Global collaborations, for example the Human Cell Atlas project, further make profiling millions of cells possible^1^. However, this trend of increasing data generation also introduces the challenge of data integration. Deep-learning-based approaches provide many solutions to integrate single-cell datasets^2-5^. Additionally, their availability to a wider research community is still limited due to the computational cost. Besides the need to reduce computational burden, we also face another challenge of standardizing data annotation. Different research groups have their own practices for cell-type annotation. The same cellular system profiled by different research groups could have different cell-type annotations. For example, the human heart atlas study defined 9 major cell types and 27 sub-types, while a similar atlas-level study defined 17 cell-types for the cardiovascular system^6,7^. Without the standardization of cell-type annotation, it is hard to establish agreement for integrative analyses. This is also a pressing issue for integrating COVID-19 related single-cell transcriptomics datasets that have been generated by researchers across the globe to understand the SARS-CoV-2 disease mechanism^8,9^.

To address these issues in integrative analysis of scRNA-seq data, we propose a fast, model-free method for standardization and integration of cell-type annotation. Our method relies on using putative cell-type markers from reference data to uniformly annotate query datasets, as putative cell-type markers are reliable indicators and should hold a constant truth across different studies. Because of its simplicity, our method can easily accommodate annotation for millions of cells using limited computational resources.

## Results

### Development of the MASI method

In our previous study, we found that converting the gene expression matrix to cell-type score matrix through a scoring method (PlinerScore^10^) based on cell-type markers in PanglaoDB^11^ can be used for integrative cell-type annotation^12^. Here, we first wanted to address the impact of different processing steps on cell-type annotation and batch-mixing. For this, we selected 4 batch-involved datasets of 4 tissues and tested 10 different processing pipelines for revealing cell-type separation, and batch-mixing (Supplementary Fig. 1). The #1 pipeline is the most basic data processing for scRNA-seq analysis, which does not take batch information into consideration for calculating the highly variable genes. The #2 pipeline differs from #1 in terms of identifying highly variable genes (HVG) by batch and only including shared HVGs in downstream processing. For #3 and #4 pipelines, we introduced cell-type markers that are obtained from either PanglaoDB or a specific reference data, and we only included those marker genes in the downstream analyses^11^. For #5 and #6 pipelines, we further converted the gene expression matrix to a raw cell-type score matrix containing cell-types in PanglaoDB or in the specific reference data. In pipeline #7 and #8, we added a transformation process (PlinerScore) as proposed by Pliner *et al*.^10^, before converting the gene expression matrix to a cell-type score matrix. #9 pipeline is a combination process of #2 and #8 pipelines. Deep-learning-based batch correction methods demonstrated a considerable success to integrative analysis of scRNA-seq data, and we noticed that the frequent practice across these methods is use of batch normalization layer and non-linear activation layer, which splits the whole dataset into multiple mini-batches, standardizes cells in each batch, and transforms the outcome with a non-linear activation function^2,4,5,13,14^. This batch normalization and non-linear activation process do not require weight training, and we included it into the #10 pipeline. These 10 pipelines were evaluated in terms of how well these pipelines preserve cell-type structure while mixing batches (Supplementary Fig. 2). We defined cell-type silhouette score to quantify how the processing pipeline reveals cell-type structure and batch entropy mixing score to evaluate how well batches are mixed. Based on our benchmark, we observed that pipelines that use conversion using cell-type markers either obtained from PanglaoDB or from a specific reference data largely mixed different datasets better, and revealed cell-type structure (pipeline #5, #6, #7, and #8), while calling HVG by batch (pipeline #2) and using cell-type markers (pipeline #3 and #4) alone resulted in a lower batch entropy mixing score. We also noticed that the pipelines with PlinerScore (pipeline #7 and #8) had a slight improvement from the raw cell-type score pipelines (pipeline #5 and #6). Both pipeline #9 and #10 have a higher batch entropy mixing score, but a lower cell-type silhouette score. For a good balance of cell-type silhouette and batch entropy mixing, we selected #8 as the processing pipeline for mapping cell-type labels for a query dataset when a reference is available (Supplementary Fig. 2).

### Workflow of MASI for integrative analysis

To annotate cell-types for a query dataset based on a fully annotated reference dataset, we propose a novel MASI workflow. MASI identifies cell-type marker genes from the reference dataset, processes data with the pipeline #8, annotates cell types via MACA^12^, and performs other downstream integrative analyses (Fig. 1a). Briefly, MACA is a marker-based cell-type annotation tool that converts a cell by gene matrix to cell by cell-type matrix, yielding a cell-type label for each identified cluster. The first step of the MASI workflow is to identify marker genes for each cell type via differential expression (DE) tests if author-verified markers are not available. To select the DE method that facilitates accurate cell-type annotation through MACA, we benchmarked 12 DE tests, including common DE tests implemented in Scanpy^15^ and Seurat^16^, and two newly proposed methods COSG^17^ and Cepo^18^. For the 4 benchmark datasets, we found that marker genes obtained from these 12 DE tests have varying performances in terms of predicting cell types using MACA (Supplementary Fig. 3). This is consistent with results shown in other benchmark studies on DE tests^19-21^, where none of single DE tests can faithfully identify reliable cell-type markers for all single-cell data. To account for influence by single DE tests, we decided to construct ranked cell-type markers via an ensemble approach (Fig. 1b and Method section for rank aggregation). Additionally, to accommodate large-scale scRNA-seq data, we refactored the MACA annotation workflow in a parallel manner by splitting data into multiple batches and distributing annotation onto multiple CPU cores (Fig. 1c). This enables MACA to perform integrative analysis for large-scale scRNA-seq with limited computational resources.

**Fig. 1:**
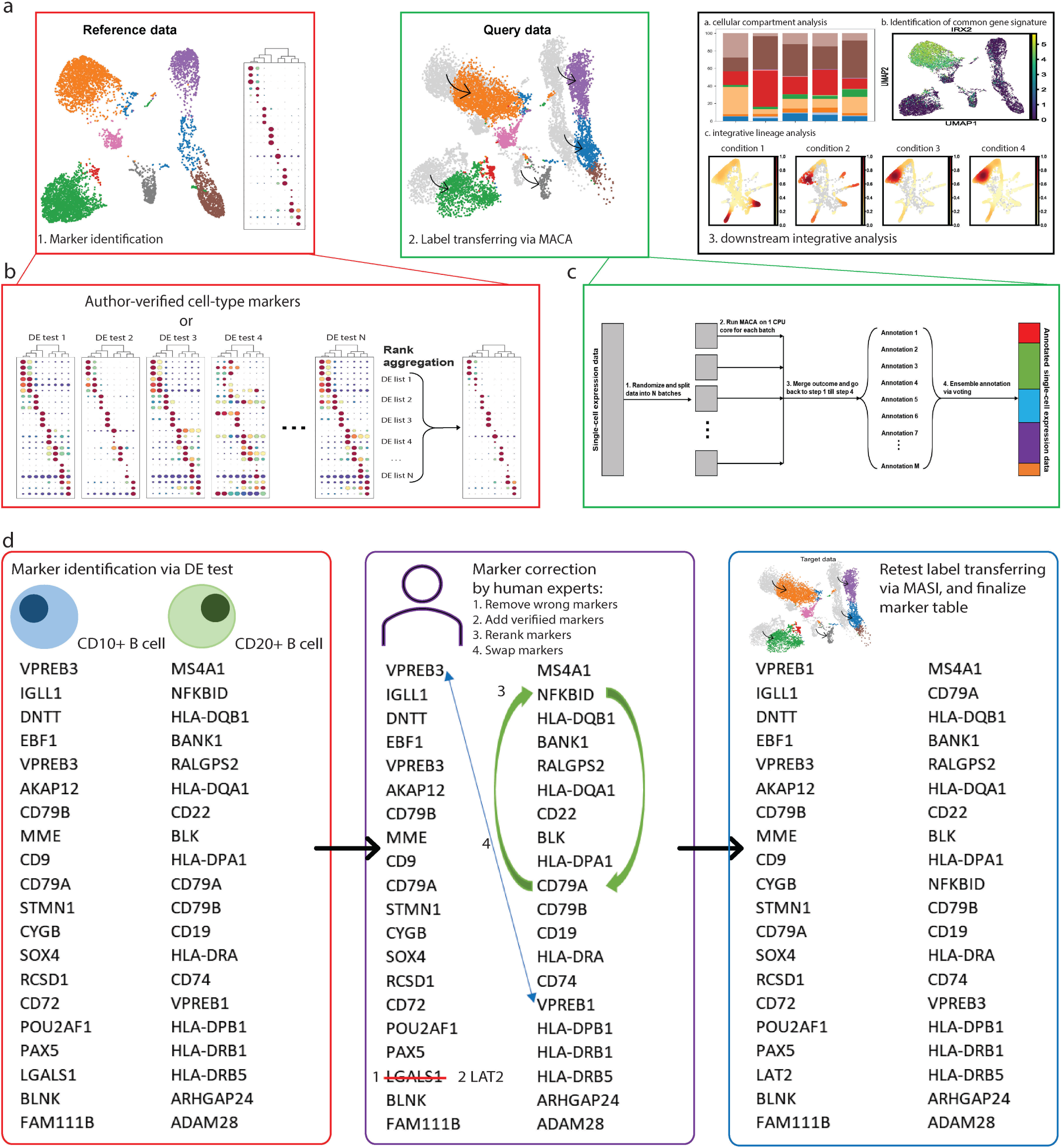
Integrative annotation pipeline through MASI. a, A workflow of integrative annotation through MASI, including marker identification from reference data, label transferring by MACA, and downstream integrative analyses. b, Ensemble approach to identify robust cell-type markers from reference data. N DE test outcomes are aggregated to get final ranked marker list. c, Parallel computation for fast annotation in order to accommodate large-scale scRNA-seq data. d, Suggested actions for improvement of label transferring. Human expert can correct wrong markers, adjust marker ranking, and so on, in order to improve annotation accuracy by MASI.

### Building a marker collection for standardized cell-type annotations

Using the ensemble approach (Fig. 1b) to automatically identify markers for cell types across species and tissues, we built a marker collection for cell-type annotation. In addition, we also added author-verified marker tables into our collection where available. This marker collection is deposited at the MASI GitHub. Additionally, our marker collection is customizable, where users can add, delete, or readjust marker gene ranking (Fig. 1d). With this marker gene collection, we could apply MASI for integrative analysis of scRNA-seq datasets in different scenarios. In the following sections, we benchmark MASI and other comparable methods for the task of cell-type annotation and data integration in terms of reliability, speed, and accuracy.

### Benchmarking cell-type annotation and data integration

We benchmark MASI and other selected methods using the 4 mixed-batch datasets that include a human pancreas data across 5 scRNA-seq platforms^22-26^, human hematopoietic data across 4 studies^27-30^, human heart atlas^6^, and mouse brain data across 4 studies^31-34^. The human heart atlas data were collected from two institutes and covered single-cell, single-nuclei, and CD45+-enriched data. We selected linear and non-linear support vector machine (SVM) classifiers as supervised methods for benchmarking as other studies have previously demonstrated that SVM outperformed most sophisticated cell-type^35^. scNym^4^ and scArches^5^ are semi-supervised deep learning methods for cell-type annotation and data integration, and we included these two methods in our benchmark. For a fair comparison, our benchmark study was performed on a local workstation with 64GB memory and Nvidia Quadro RTX 6000 as GPU support. Of note, both scNym and scArches use GPU to speed up computation, while MASI will not use GPU for computing. We first focused on how well mapping cell-type labels from reference data to query data is done by these methods. We used macro F1 and overall accuracy to quantify the performance of these methods in terms of how accurate annotation is for each cell type and how accurate annotation is for the overall dataset. We found that all methods have similar performance in terms of overall accuracy, but MASI has higher macro F1 scores across all datasets in our benchmark (Fig. 2a). This suggests an advantage of MASI in annotating non-major cell types, considering most single cell data are class imbalanced.

**Fig. 2:**
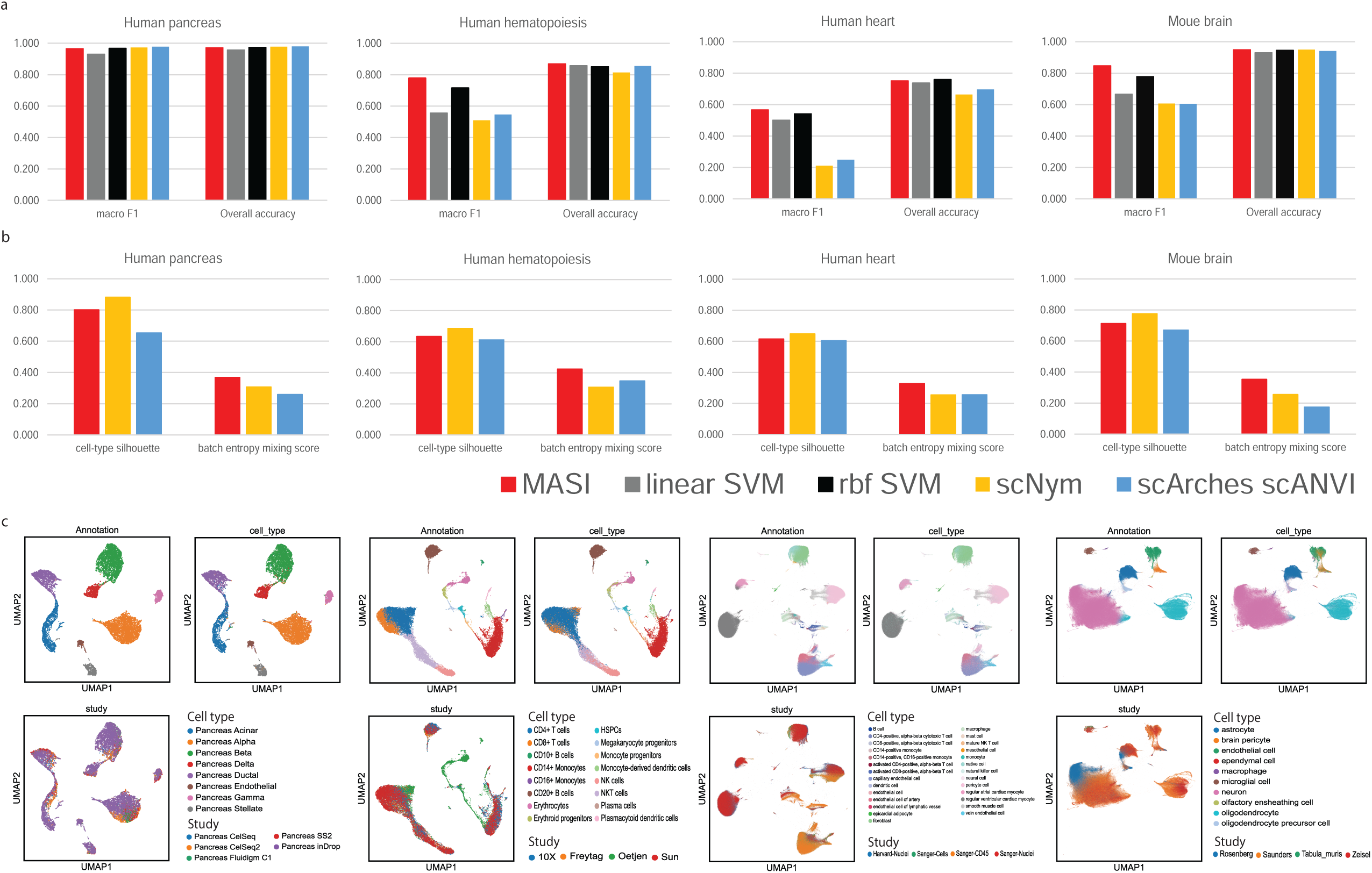
Batch correction and label transferring benchmarks. a, Comparison of label transferring for MASI, supervised, and semi-supervised methods. ACC: overall accuracy. Macro F1 is average of F1 scores per cell type. A higher score in both metrics suggests better cell-type prediction. b, Comparison of batch correction for MASI, scNym, and scArches scANVI. Cell-type silhouette score measures how well the integrated representation by these methods preserves cell-type variation, while batch entropy mixing score measures how well the same cell type from different batches is mixed. c, Visualization of MASI integration through UMAP. Cells are colored according to MASI-reported annotation (top left), author-reported annotation (top right), and batch id (bottom left).

Next, we evaluated how well the representations learned by these methods capture the cell-type variations while mixing batches, using cell-type silhouette score and batch entropy mixing score mentioned above. Here, MASI demonstrated a good balance between capturing cell-type variation and batch mixing (Fig. 2b). We found all 3 methods, MASI, scNym, and scArches, achieved the same purpose of mixing data from diverse sources (Fig. 2c and Supplementary Fig. 4). However, we observed that representations learned by scNym and scArches captured weaker spearman correlations among different cell-types, while the cell-type score representation of MASI preserved the cellular correlation, especially in human hematopoiesis (Supplementary Fig. 5).

### Dependence on choice of reference dataset and clustering resolution

Given the high dependence on reference data for cell-type marker identification, MASI will not be able to annotate cell types in query data that have not been seen in reference data. However, it is still worth answering if a cell-type score matrix constructed using the reference data can preserve cell-type structure for query data, even though query data contains unseen cell types. To understand the impact of choice of reference dataset on the cell-type annotation in the query dataset, we swapped reference data from Oetjen *et al*. to 10x Genomics data in the human hematopoietic benchmark dataset. The 10x Genomics data has only 12 cell types, while Oetjen *et al*. identified16 cell types in their original report. Thus, the query data would contain 4 extra unseen cell types. We performed marker gene identification and transformed the gene expression matrix to cell-type score matrix as above. We observed that the 12-dimension cell-type score matrix built upon the 10x Genomics dataset as reference can reveal cell-type structure for the Oetjen *et al*. data that had 16 author-reported major cell types in total^27^. However, as erythrocytes and erythroid progenitor cell-types are not present in the reference, MASI mislabeled them as CD14+ monocytes and HSPCs, respectively (Supplementary Fig. 6). We next asked if we could identify subtypes from MASI-reported cell types to match the author-reported annotation resolution. Here, we used SCCAF, a computational method that was previously proposed for the identification of putative cell types through a machine learning approach^36^. The concept behind this machine learning is: if the clustering resolution reflects the number of true cell types within the data, a machine learning classifier can achieve a high accuracy with the clustering label. Thus, we applied SCCAF to identify potential subtypes for each major cell type identified by MASI. We evaluated how these three approaches, MASI annotation, SCCAF identification, and SCCAF+MASI annotation respectively, revealed a similar annotation resolution by calculating ARI and NMI against the author’s annotation. We found that SCCAF+MASI annotation matches the author’s annotation resolution more than MASI annotation and SCCAF identification alone (Supplementary Fig. 6). To summarize cell-type identification, we conclude that the choice of reference data is critical to the performance of MASI. Moreover, to unravel potential subtypes, users can combine SCCAF and MASI to reach a finer annotation.

### Annotation of spatial transcriptomics data with MASI

Next, we used MASI to map cell type labels from scRNA-seq data to sequencing-based spatial transcriptomics data. We tested this on spatial hippocampus data profiled by Slide-seqV2, since Slide-seqV2 reaches a higher resolution of spatial profiling than 10X Visium^37^. Integrating Slide-seqV2 with scRNA-seq further suggests a potential application of MASI in spatial transcriptomic analysis (Supplementary Fig. 7a). MASI was able to assign cell type labels to the mouse hippocampus Slide-seqV2 data (Supplementary Fig. 7b). Spatial expression patterns of marker genes for 5 distinct cell types also match with their cell locations in space (Supplementary Fig. 7c and d).

### Integrative temporal analysis using cell-type score matrix

An advantage of using cell-type scores as features is that it condenses biological information from high-dimension gene feature space into a lower dimension cell-type feature space. Meanwhile, we also showed above that our approach is useful for mixing batches coming from different studies. Next, we utilized these cell-type features to integrate multi-batch temporal scRNA-Seq datasets for lineage analysis. For this, we selected three datasets for lineage analysis: 1) human peripheral blood mononuclear cell (PBMC) data of patients with Kawasaki disease obtained before and after IVIG (intravenous immunoglobulin) treatment^38^, 2) mouse brain lineage tracing at different time points^39^, and 3) zebrafish embryo from two studies that cover 13 major developmental stages^40,41^.

Both human hematopoiesis and mouse brain lineage tracing studies were in multi-condition design, and we were able to obtain cell-type markers from an externally annotated 10X Genomics PBMC data^30^, while author-verified cell-type markers from the original report were used for the mouse brain lineage tracking study^39^. We constructed an integrative lineage map with cell-type score matrices and visualized population density and cell-type score (Fig. 3a and Supplementary Fig. 8). Using cell-type scores to interpret data, we were able to identify lineage changes, which is consistent with the original report. For example, we observed decreased B1 B-cell and CD16+ monocyte lineages as well as restored plasma cell and CD4+ T native lineages after IVIG treatment for acute Kawasaki disease patients (Fig. 3a). In mouse brain lineage tracing study, the integrative lineage map showed that mitotic progenitor cells injected at E10.5 time point tend to differentiate to astrocyte and OPC (oligodendrocyte precursor cell), while the lineage specification may shift to neuron at 14.5 time point (Supplementary Fig. 8). This is also consistent with the original findings.

**Fig. 3:**
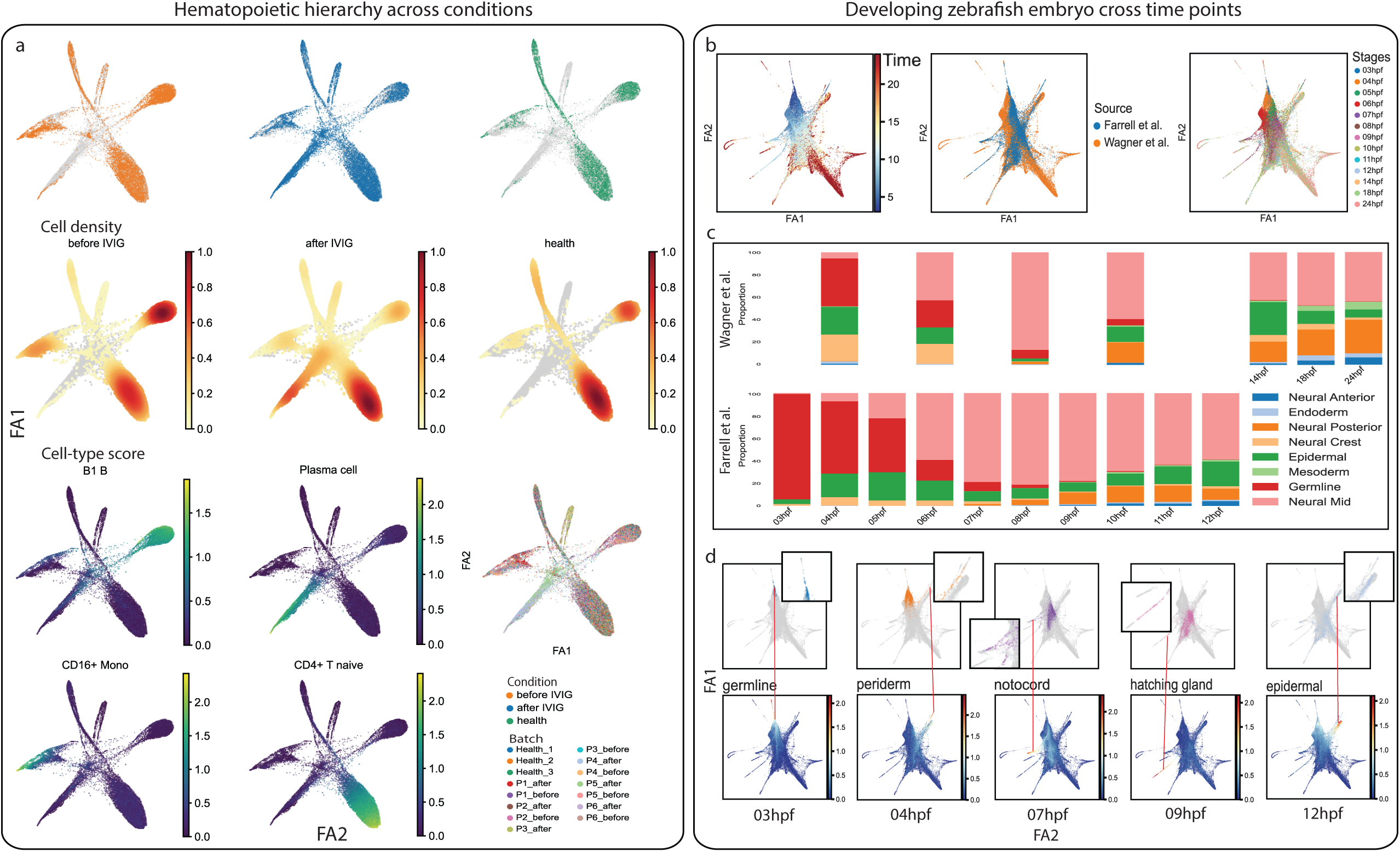
Integrative lineage analysis using cell-type score matrix. a, Integrative lineage analysis for multi-condition human hematopoiesis study. Cell density, cell-type score, and batch id for human hematopoiesis samples under different conditions are visualized separately through the first two ForceAtlas2. b, integrative lineage analysis for two developing zebrafish embryo data. Cells are visualized through UMAP and are colored according to developmental time (left), study id (middle), and developmental stages (right). c, Components of 8 major lineages along the developmental stages in zebrafish embryo. All lineages sum up to 1 in one stage, and data from the two studies are visualized separately. d, Identification of lineage origin time. Visual investigation is conducted by matching emergence of a cell-type with the earliest developmental stage in the data.

Our integrative analysis of developing zebrafish embryos consists of data from two independent data sources that cover different time points of post fertilization. Wanger *et al*. collected cells from 7 stages including 4, 6, 8, 10, 14, 18, and 24 hpf (hours post fertilization), while Farrell *et al*. designed 12 finer stages ranging from 3 to 12 hpf. We were unable to find an external marker gene reference for the two developing zebrafish datasets. Given they were in a time-series design, we reasoned that the end-point data should contain all mature cell types. Therefore, we intrinsically selected the end-point data that has 30 cell types as reference to identify cell-type markers. In total, these 2 independent studies cover 30 cell types along the 13 developmental stages. Next, we transformed the combined gene expression matrix into a 30 cell-type score matrix and built an integrative lineage map of the developing zebrafish embryo. Because of the design differences, we manually summarized all developmental stages in 13 major stages (Fig. 3b). Instead of assigning cells to these 30 cell types, we annotated them as 8 major lineage types using MASI. The choice for these 8 major cell-types was based on the lineages observed in 24 hpf (refer to wagner et al). Markers for these 8 major lineage types were also identified from data at the 24 hpf time point. We then visualized how lineage components change along the developmental timeline (Fig. 3c). First, we found that the two studies are largely consistent. Second, we observed a decline of germline and lineage diversification along these developmental stages (Fig. 3c). We further investigated the original time point of different cell lineages based on our integrated lineage map and found that the development of germline can be retrieved back at least at the 3 hpf time-point (Fig. 3d). In Wagner *et al*., the earliest time point at which germline cells were observed is 4 hpf. However, in Farrell *et al*., the authors report that the germ layer appears before 4 hpf and that many other lineages do not separate until 4 hpf. This is consistent with our finding from the integrated lineage map, where we show that germline cells are observed at 3 hpf and are the major cell lineage component until that time point (Fig 3c and Fig. 3d). The notochord defines the longitudinal axis of the embryo and determines the orientation of the vertebral column, and our analysis suggests the notochord emerges at around 7 hpf, while both Farrell *et al*. and Wagner *et al*. showed emergence of the notochord takes place between 6hpf and 8hpf. We also observed that epidermal lineage appears at 3 hpf (Fig. 3c), consistent with Farrell *et al*. who observed this epidermal lineage at an early time point (3.3 hpf). Additionally, we observe that non-neural ectoderm separates from epidermal cells at 12 hpf in our analysis, as seen in Farrell et al. (). Taken together, these three analyses for temporal datasets using MASI shows a simple and intuitive approach for integrative lineage analysis.

### Case studies to explore data integration and standardization in large datasets

We applied MASI to three case studies consisting of 783976, 486134 and 251057 cells from heart, kidney and COVID19 datasets. The datasets consist of 27, 27, and 59 cell types, respectively.

### Case 1: Cell-type annotation and batch-mixing of human heart datasets

Tucker *et al*.^*7*^ and Litviňuková *et al*.^6^ provide two atlas level resources for human heart data at single-cell resolution. In addition, other human heart datasets are also available^42,43^. However, these studies did not use the same cell-type naming style and reported annotation at different resolutions. Litviňuková *et al*. identified 27 subtypes while Tucker *et al*., (17 subtypes), Wang *et al*. (5 cell types), and Cui *et al*. (9 cell types) reported different numbers of cell-types in their datasets^6,7,42,43^. We think uniform annotation of cell-type labels and batch-mixing of these datasets can yield insights into common themes and inter-human variability across these datasets. Since the human heart atlas data revealed the greatest number of subtypes, we chose human heart atlas data as reference. Because cell-type naming and annotation resolution vary among these studies, we changed to use ARI and NMI for evaluation. We found all methods compared here have similar performance for mapping cell-type labels to Tucker *et al*., but MASI shows better outcome than the other 4 methods in both Wang *et al*. and Cui *et al*. (Fig. 4a). Moreover, both Wang *et al*. and Cui *et al*. have distinct difference in the number of cells profiled and both have a lower annotation resolution than Litviňuková *et al*. and Tucker *et al*. Relying on a greater resolution of Litviňuková *et al*. data, we were able to level up annotation for the other 3 studies (Fig. 4b). We visualized integration via MASI and observed no distinct batch differences (Fig. 4c). We noticed MASI annotated a number of cells in Tucker *et al*. as fibroblast while the author-reported annotation for these cells includes cardiomyocyte, endothelium, and neural cells (Fig. 5d). We looked deeper into these cells and examined expressions of marker genes for fibroblast, endothelium, and neural cells and found that the disagreement between MASI-reported and author-reported annotation is likely due to background mRNA from fibroblasts (Supplementary Fig. 10). With MASI, we identified pericyte in Wang *et al*. and natural killer cell in Cui *et al*., which were not reported by authors (Fig. 5d and Supplementary Fig. 11).

**Fig. 4:**
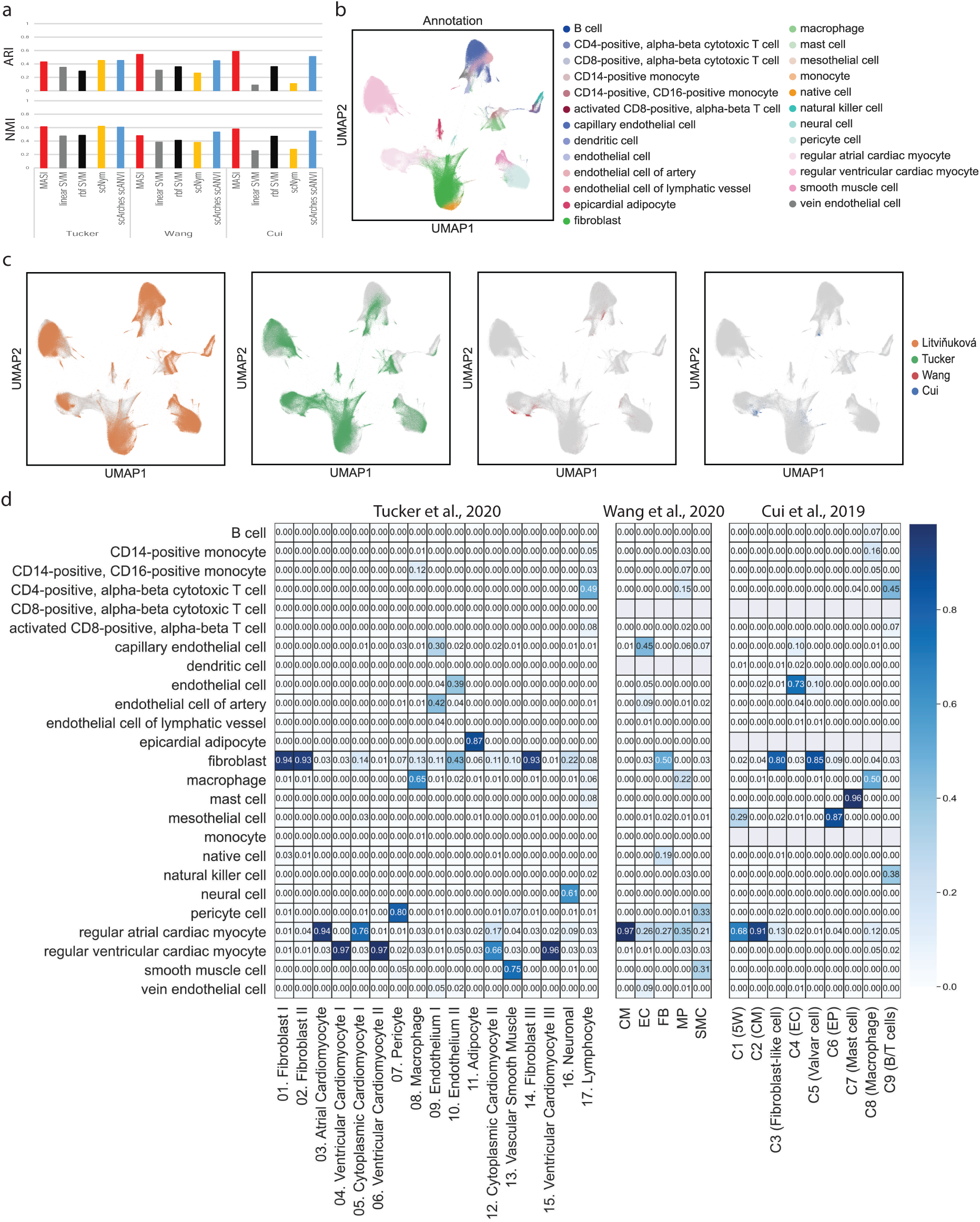
Transferring human heart atlas for integration of single-cell human heart across research groups. a, Comparison of label transferring for MASI, supervised, and semi-supervised methods. ARI and NMI are calculated by comparing method-reported annotation with author-reported annotation in a study-wise manner. b, Visualization of the integrative annotation by MASI. Cells are colored according to MASI-reported cell-type annotation. c, Visualization of integration by MASI. Cells are colored according to study id. d, confusion matrix of MASI-reported annotation against author-reported annotation. Confusion matrix is normalized to have column sum as 1. Row names use the naming style of human heart atlas, and column names remain the original naming styles of Tucker et al, Wang et al, and Cui et al data.

**Fig. 5:**
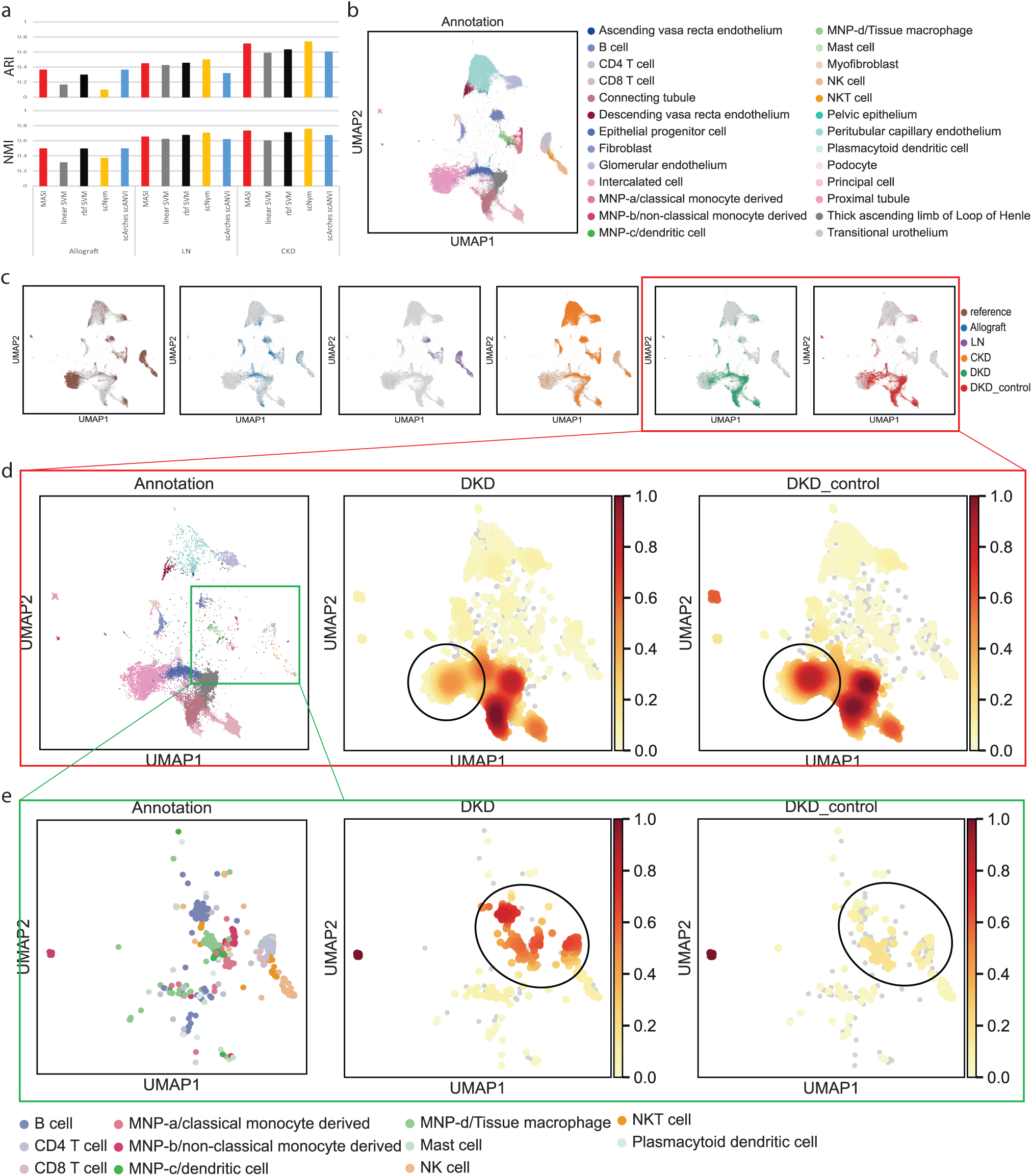
Transferring human kidney atlas for integration of single-cell human kidney across conditions. a, Comparison of label transferring for MASI, supervised, and semi-supervised methods. ARI and NMI are calculated by comparing method-reported annotation with author-reported annotation in a study-wise manner. b, Visualization of the integrative annotation by MASI. Cells are colored according to MASI-reported cell-type annotation. c, Visualization of integration by MASI. Cells are colored according to study id. d, Population densities in DKD and control samples. Cell type annotation is shown on the left panel. Cell type population densities of DKD and control samples are presented separately to highlight difference of cell type populations. e, Population densities of immune cells in DKD and control samples. Cell type annotation is shown on the left panel. Cell type population densities of DKD and control samples are presented separately to highlight difference of cell type populations.

### Case 2: Using human kidney atlas for integration of single-cell human kidney across multiple conditions

The first human kidney atlas profiled 27 distinct cell types in mature kidney^44^. This atlas provides a good reference to study cellular irregularities in kidney diseases. So far, independent single-cell studies have been conducted to reveal mechanisms in different kidney diseases^45-48^. An approach that can provide an integrative view for multiple kidney diseases may further add an insight into how cellular irregularities vary among different kidney diseases. We used kidney atlas data as reference and mapped cell-type labels to human kidney data that were collected under different conditions, including CKD (chronic kidney disease) and DKD (diabetic kidney disease). Benchmarking in this task showed MASI has better agreement with author-reported annotations with consistency (NMI values of 0.49, 0.648, and 0.728, respectively) (Fig. 5a). Overall mapping, cell type standardization and batch-mixing results are shown in Fig. 5b and 5c. Next, we focused on the human DKD data, which came with its control set. Population density map suggested decrease of proximal tubule (Fig. 5d) and increase of immune cells (Fig. 5e), consistent with an increase of immune response identified in DKD patients^46^.

### Case 3: transferring human lung atlas for integration of single-cell COVID19 data across participants

Our third MASI application is transferring knowledge learned from human lung atlas to understand the global COVID19 pandemic at cellular level among healthy and COVID19 participants. The human lung atlas data served as reference data with 59 identified subtypes^49^. Using this annotation, we aimed to annotate 80 COVID19 samples collected from nasal swabs (58 participants and 32818 cells) and airways across different individuals (22 participants and 143168 cells)^8,9^. These COVID19 data included both negative (21 participants) and positive samples (59 participants) from multiple centers. Due to cell-type annotation and resolution differences, we cannot directly compare cellular differences between healthy and COVID19 participants. We used MASI to annotate the COVID19 data to match the annotation resolution of human lung atlas. Again, we benchmarked MASI with two SVM classifiers, scNym, and scArches, using ARI and NMI as evaluation metrics. We found MASI has greater agreement with author-reported annotations for all COVID19 data than the other 4 methods (Fig. 6a). Since cell-type annotations for all participants were leveled up to the same resolution, we were able to directly compare the cellular differences between healthy and COVID19 participants (Supplementary Fig. 12). We observed distinct cellular components between healthy and COVID19 groups, and the distinct cellular component is consistent across participants within the same group (Fig. 6b). Then, we quantified the changes of cellular composition for all cell types and found an increase in the proportion of Goblet cells and a decrease in ciliated cell proportions in the COVID19 group. This discovery may explain other investigations of SARS-CoV-2 virus targeting ciliated cells via *ACE2*^*50,51*^.

**Fig. 6:**
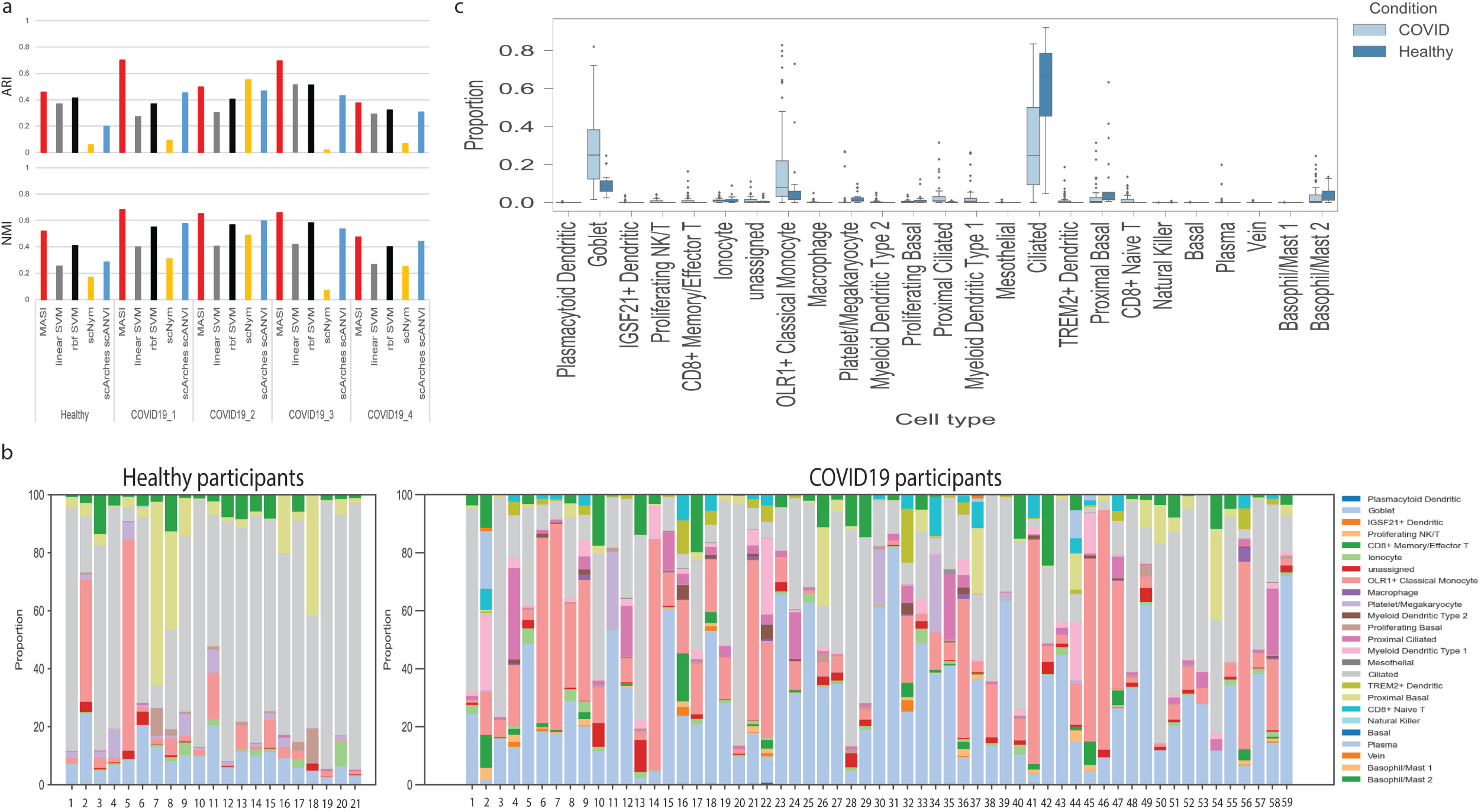
Transferring human lung atlas for integration of single-cell COVID19 data across individuals. a, Comparison of label transferring for MASI, supervised, and semi-supervised methods. ARI and NMI are calculated by comparing method-reported annotation with author-reported annotation in a study-wise manner. b, Cellular components of healthy and COVID19 participants. Each column represents one individual. c, Cellular component comparison of healthy and COVID19 participants.

### MASI is fast and can accommodate annotation for large scale single-cell data

Taken together, we show that MASI can quickly annotate large-scale scRNA-seq data. Our runtime test showed runtime of model-based methods increases dramatically once the data scale up (Supplementary Fig. 13). For an extreme test, we performed integration of mouse brain data (nearly 1 million cells) by MASI on a personal laptop, with 16GB memory and no GPU support. Without sacrificing annotation accuracy, MASI can scale up to accommodate label transferring for 1 million cells (Supplementary Table 1).

## Discussion

Here, we present MASI, a new tool to quickly and accurately annotate single-cell datasets based on marker genes obtained from a reference dataset. We show that MASI can also be used for batch-mixing and serve as a data integration method for single-cell transcriptomics data. We benchmarked MASI with supervised and semi-supervised methods, and our results show that performance of MASI is comparable or slightly better than other tested methods based on the datasets used in this study. We also showed that cell-type scores can be used as features for integrative lineage analysis and demonstrated its intuitive interpretability. Finally, we showed the utility of MASI in three different case studies of data integration covering different research groups, biological conditions, and surveyed participants. Like other supervised and semi-supervised methods replying on reference data, accurate annotation via MASI is also dependent on the quality of reference data. Thus, the choice and resolution of the reference are critical to downstream analysis. Therefore, we recommend users to select reference data that has annotation resolution compatible with their downstream investigations. If query data has unseen cell types not in reference, MASI in combination with SCCAF can be used to identify subtypes within major cell-types. Additionally, we showed that MASI can also be applied for cell-type prediction in spatial transcriptomics datasets using comparable single-cell transcriptomics datasets as reference.

There are many well-established integration methods available to address batch effects in scRNA-seq datasets, for example Seurat, Harmony, and LIGER^16,52,53^. Additionally, some deep learning-based methods such as HDMC and CarDEC are also available^54,55^. In this study, we rigorously tested cell-type score-based integration via MASI across various single-cell platforms, cytoplasm/nuclei, research groups, conditions, and individuals. Our analyses suggest marker-based feature engineering can be useful for reference-based cell-type annotation, batch-mixing, and data integration. We also demonstrate that integration via MASI preserves biological information for lineage analysis with 3 different examples.

Overall, MASI is easy to set up and requires limited computation resources to run. It can be used for reference-based cell-type annotation and batch-mixing, which could facilitate quick hypothesis-driven exploration of diverse datasets obtained from different labs. Moreover, the democratization of single-cell transcriptomics data (larger cellular output with lower cost) could empower researchers even with limited computational resources to investigate millions of single cells among diverse biological systems^56^.

## Methods

### Data preprocessing

Raw gene expression count data were ‘LogNormalized’, which divides the total count in that cell and multiplies it by a scale factor of 10 000 (in all our analyses), followed by log-transformation to get the normalized expression matrix. For implementing MASI, we skipped the step of calling highly variable genes, because only the identified marker genes were used for integrative annotation. For training scNym and scArches, we used top 5000 highly variable genes by batch, which were calculated using function “pp.highly_variable_genes” in Scanpy^15^

### Marker rank aggregation

We considered two ensemble marker ranking schemes. In the first scheme, the top 20 marker genes from each DE test were compiled together. For the second scheme, only statistically significant marker genes based on the p-values corrected for multiple hypothesis correction were considered. In the first scheme, we searched the consensus ranking via robust rank aggregation^57^. In the second scheme, rank aggregation was done through Lancaster combination^58^.

### Weighing markers

When data to be annotated contains distinct cell types and cell types do not share marker genes, we reasoned that weighing markers would not influence the final annotation by MASI. However, this can be beneficial to distinguish cell subtypes that share common markers, for example subtype T cells. We used a simple weighing strategy that returned good label transferring. Given *N* markers for cell type *A*, the 1^st^ marker in this ranked list will contribute 100% of its expression to the cell-type score of *A*, while the *N*^*th*^ marker only contributes 50% of its expression. For the *i*^*th*^ marker in the rest, we form this discount calculation 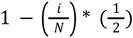 to get their weights in cell-type *A*. Beside the weighing strategy above, other weighing strategies, including Rank Order Centroid and Ratio method, can also be considered for customization.

### Converting gene expression matrix to cell-type score matrix

Cell-type score for a given cell-type *A* with *N* expressed markers is calculated by summing up the expression of all N markers with consideration of weighing markers as above. This is defined as the raw cell-type score. From this, the PlinerScore is calculated by adding a TF-IDF transformation and suppressing expression values of a marker gene to zeros if they are below the *X*-percentile of expression values across all cells before the raw cell-type score conversion. The default value for PlinerScore threshold is 0.25 as the percentile threshold^10^.

### Classification by linear and non-linear SVM

Both linear and non-linear SVM classifiers can be impacted by feature selection. As a benchmark reported, linear and non-linear SVM can have varying prediction accuracies for scRNA-seq data, when different feature selection processes were applied^59^. Nevertheless, using more discriminative features should improve the accuracy of these two supervised models. Instead of using highly variable genes and PCA-reduced features, we used the same cell-type markers that were used for MASI to train both linear and non-linear SVM classifiers. This is primarily because we observed cell-type markers have a good balance between cell-type preservation and batch-effect removal, compared to both highly variable genes and PCA-reduced features, as shown in Supplementary Fig. 2.

### Transfer learning through MASI

Once cell-type markers are identified, Mapping cell-type labels to query data is performed using MACA^12^. Briefly, for each cell, MACA generates two labels - a per-cell cell-type Label 1 and group-based clustering Label 2. Then, MACA maps clustering Label 2 to cell-type Label 1 to get the overall cell-type annotation. In our previous study, we used different clustering parameters to generate multiple Label 2s, for the purpose of reproducibility^12^. In this study, we also ran Louvain community detection with a range of clustering parameters to get multiple clustering Label 2s. These include clustering resolution 3, 5, 7 with 5, 10, 15 as neighborhood sizes to over-cluster cells. With multiple clustering Label 2s, we were able to map them to Label 1 and get a more reproducible ensembled cell-type annotation. To accommodate for large-scale scRNA-seq data, we split the whole data into N batches and ran MACA with one batch per CPU core.

### Transfer learning using scNym and scArches

Both scNym and scArches are deep-learning-based transfer learning methods. Therefore, an optimal outcome for a specific data might require customized parameter tuning. However, for benchmarking, we used default pipelines of both methods for all data involved in this study. Respective tutorials can be found at their host GitHub^60,61^.

### 2D visualization using UMAP

To visualize integrations by these three methods, we used the same parameter setting for all datasets. We set up metrics “cosine” to define distance, cells within 0.1 were considered as neighbors, and minimum 15 cells form a community.

### Integrative lineage analysis

We used ForceAtlas2 with PAGA (partition-based graph abstraction) initialization to layout integrative lineage maps with cell-type scores instead of any other hidden space features, like PCA (Principal Component Analysis) representation or representation from neural network model^62,63^. To initialize PAGA, we performed Louvain community detection to assign cells as multiple meta cells^64^. We used resolution 5 for Louvain community detection in order to get enough meta cells. Once cells are laid out on the ForceAtlas2 space, we directly visualize lineage paths with cell-type scores, without clustering cells into cell types.

### Evaluation metrics

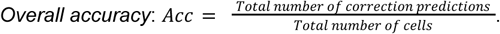

#### Macro F1

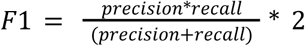. *F1* was calculated for each cell type, then *macro F*1 is the average of *F*1 scores for all cell types. Because this metric doesn’t consider class weights for imbalanced data, a higher *macro F*1 could suggest correction predictions for both dominant and non-dominant cell types.

#### Cell-type silhouette score

We first used function “sklearn.metrics.silhouette_score” in scikit-learn Python package to calculate a typical silhouette score *S*^65^. The author-reported cell type label served as the ground truth. This calculation uses the hidden space returned by integration methods with cell-type labels. Both scNym and scArches learned a 10-dimension hidden space representation by default. The lower representation by MASI depends on the number of unique cell-type labels available in the reference dataset. Next, we rescaled the score from 0 to 1 by (1 + *S*)/2 to be defined as the cell-type silhouette score. The higher the score is, the better cell-type variation is captured.

#### *Batch entropy mixing score*^66^

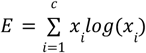. In this study, *x*_*i*_ is the proportion of cells from batch *i* in a region of the first two UMAPs, and 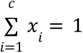. This score should quantify how well mixed cells from different batches are in a region. The same as *Cell-type silhouette score*, calculation of *Batch entropy mixing score* is based on the hidden space returned by integration methods with batch information as label. The higher the score is, the better the mixing.

#### Adjusted rand index (ARI)

The rand index (RI) measures a similarity or agreement between two clustering labels. The ARI then is defined through 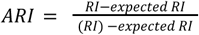. In this study, we used ARI to measure the agreement between cell-type annotation reported by a transfer learning method and the author-reported cell-type annotation.

#### Normalized mutual information (NMI)

Like ARI, NMI also qualifies the agreement between two clustering labels. It is defined as 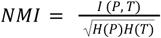. *P* and *T* are empirical categorical distributions for the predicted and real clustering, *I* is the mutual entropy, and *H* is the Shannon entropy.

## Data availability

All datasets used in this study are publicly available (Supplementary Table 2). Use of these datasets, either as reference or query data, is also specified in the Supplementary Table 2. Raw data can be found through their associated publications. Ready-to-use data are available for some datasets, and downloadable links are provided in the Supplementary Table 2.

## Code availability and reproducibility

The source code of MASI including analyses of key results in the study can be found at https://github.com/hayatlab/MASI. To reproduce results in this study, please follow tutorials deposited at (https://github.com/hayatlab/MASI/tree/main/tutorial).

## Acknowledgements

This work was supported by NIH NIGMS grant R35GM133557 to R.P.M.

## Author contributions

Y.X. and S.H. planned and designed the study. Y.X. performed the computational analysis. Y.X. and S.H. analyzed and interpreted the data and wrote the manuscript. R.P.M. and R.K. edited the manuscript and advised on data interpretation. All authors read and approved the manuscript.

