## Supplementary Fig for "Fast model-free standardization and integration of single-cell transcriptomics data"

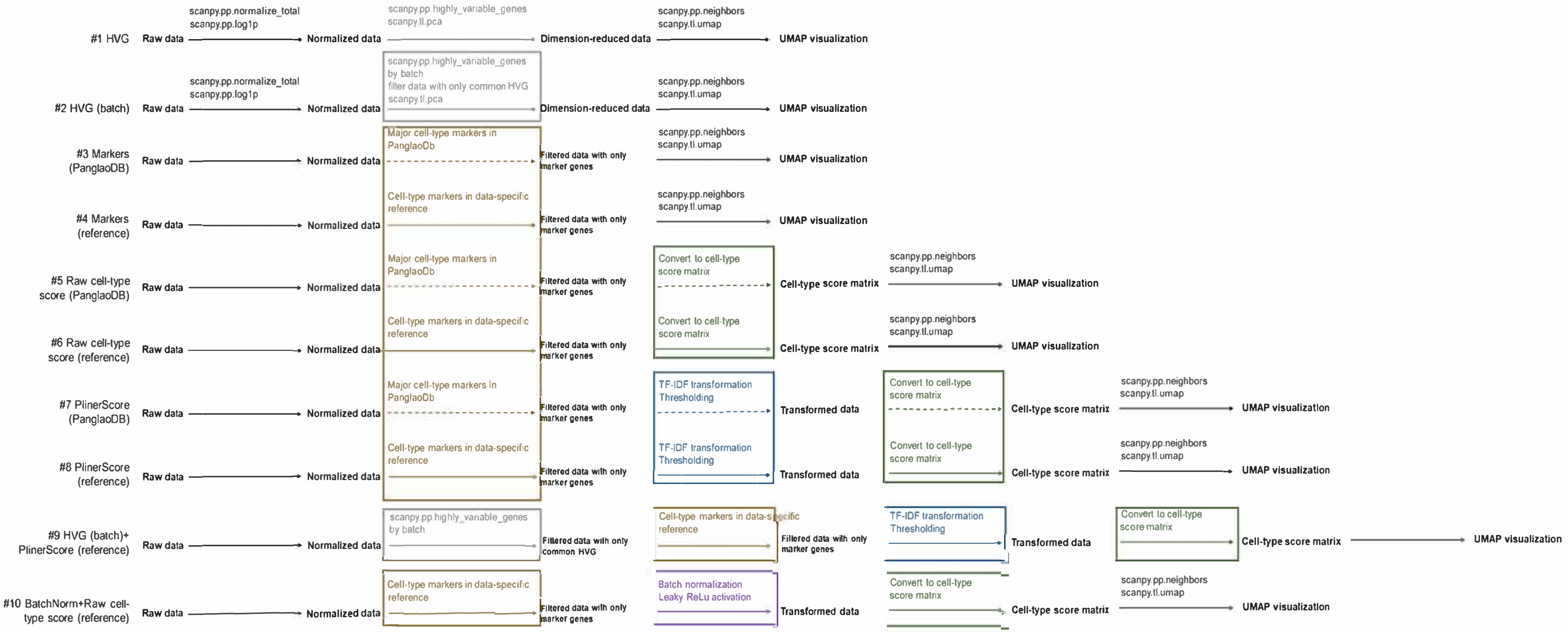


Supplementary Fig. 1

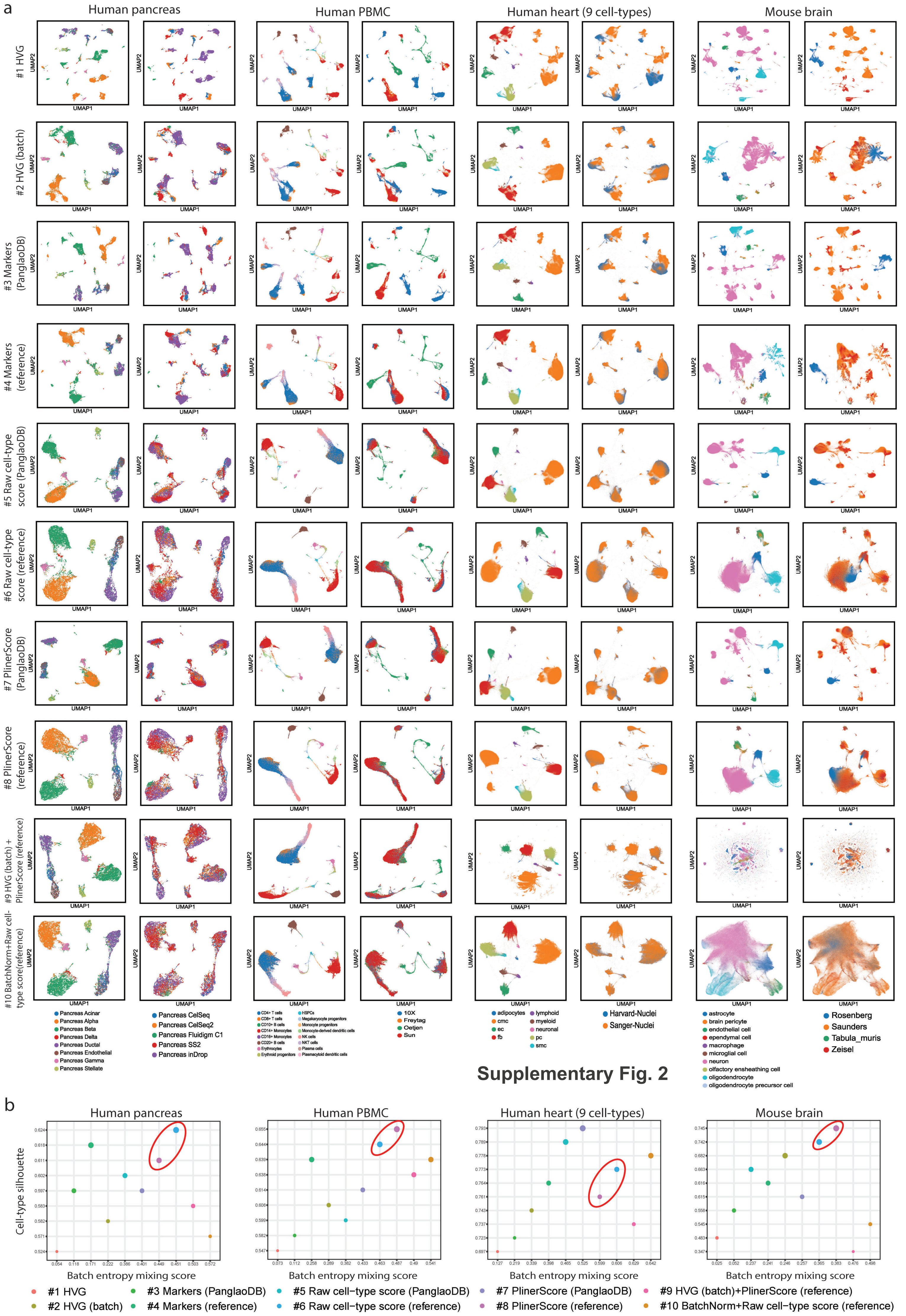


Supplementary Fig. 2

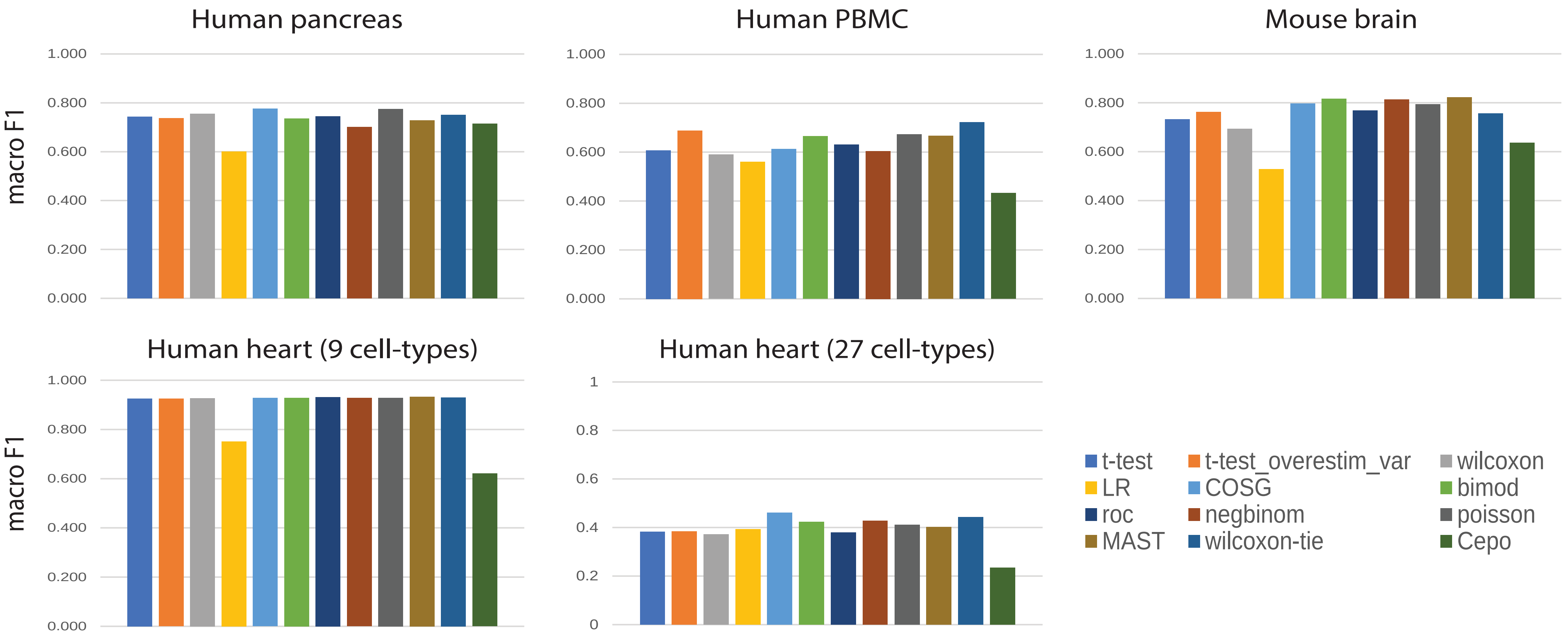


**Supplementary Fig. 3**

**Supplementary Fig. 4**

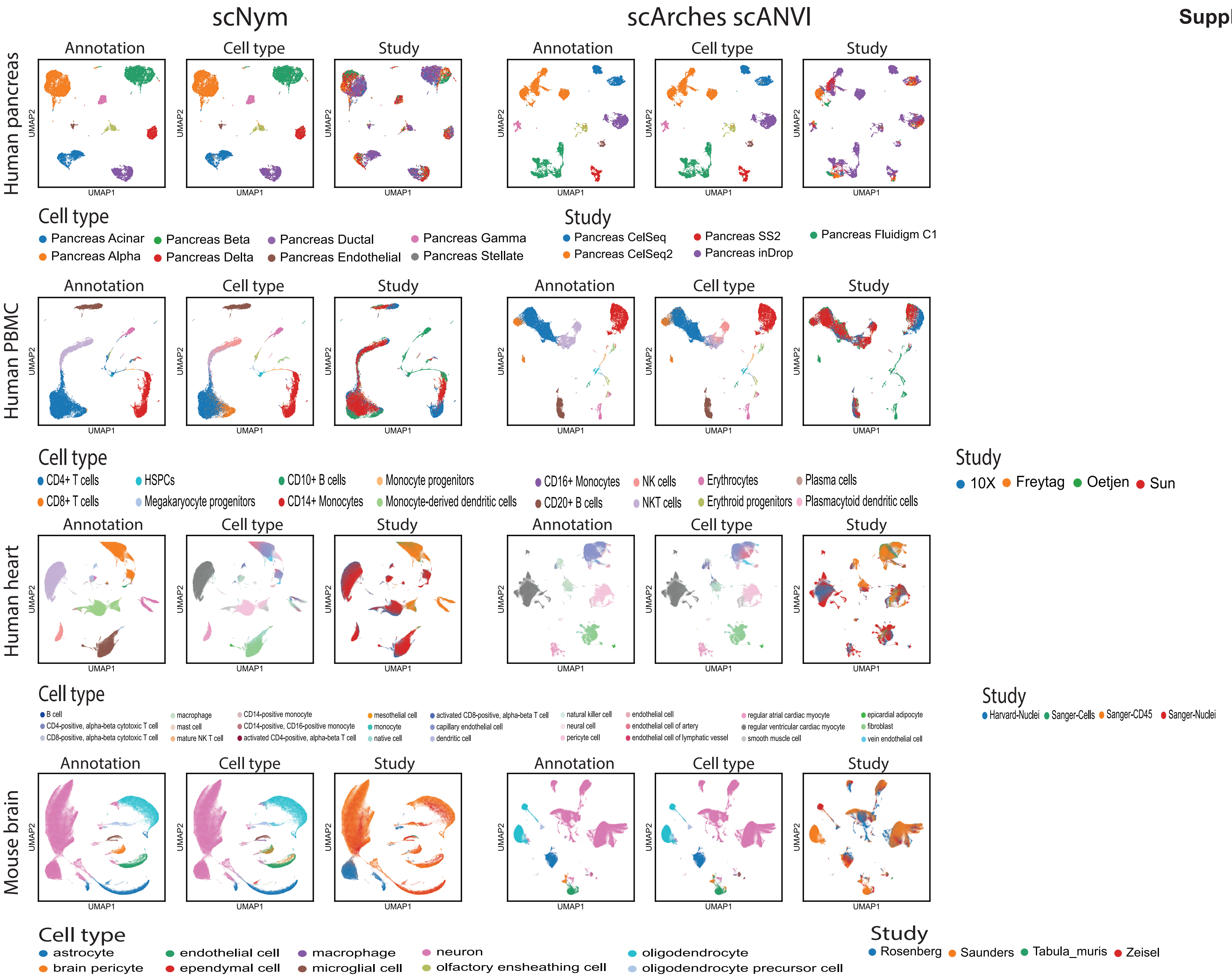

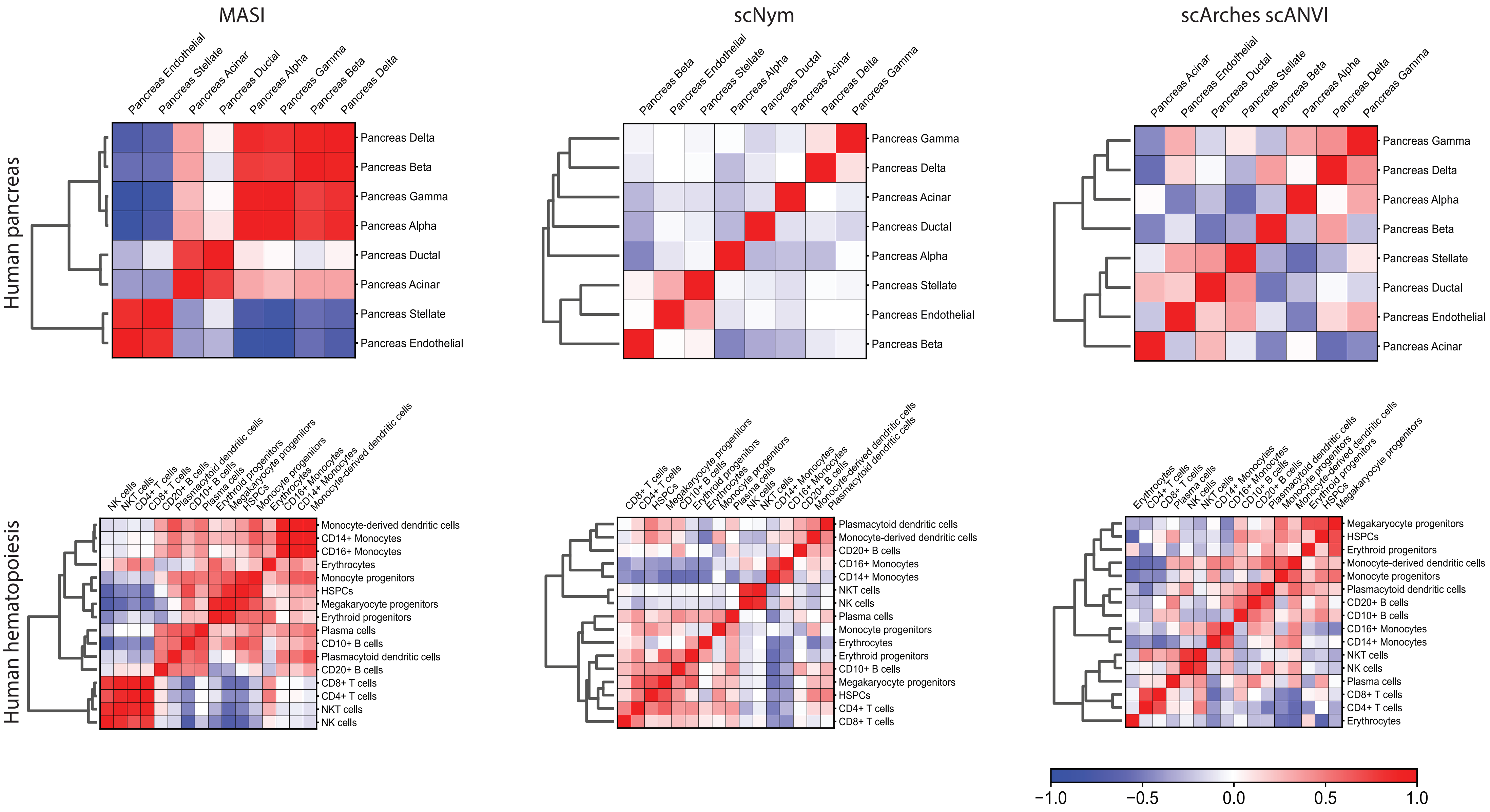


**Supplementary Fig. 5**

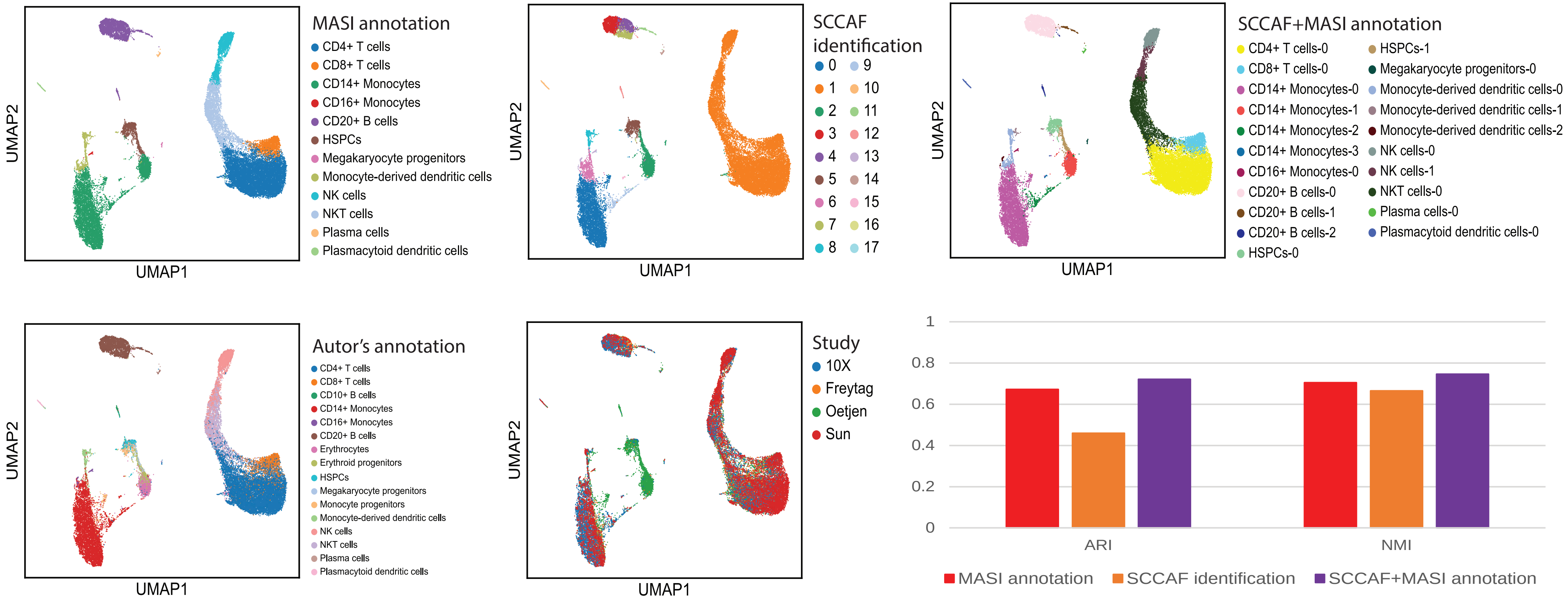


**Supplementary Fig. 6**

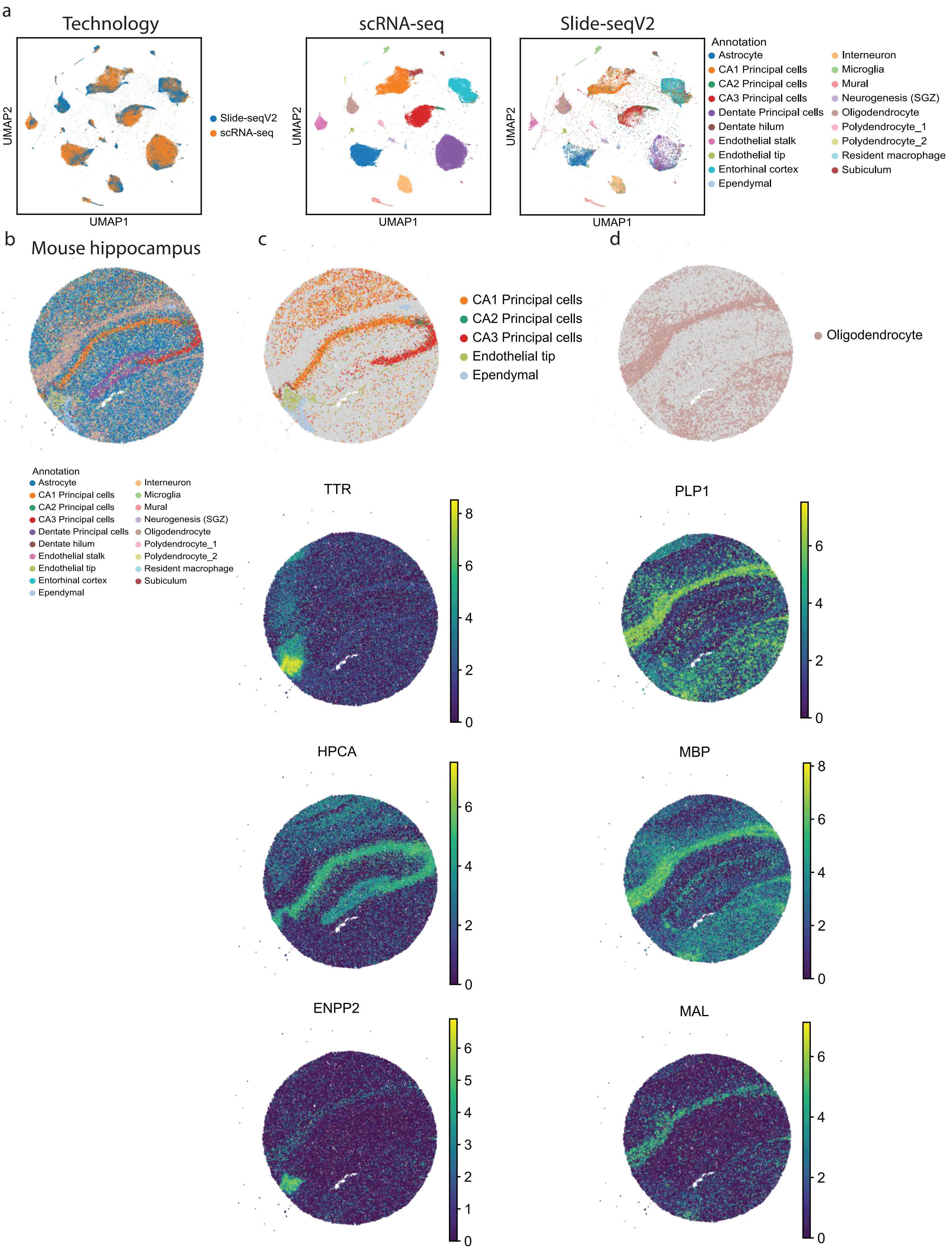


**Supplementary Fig. 7**

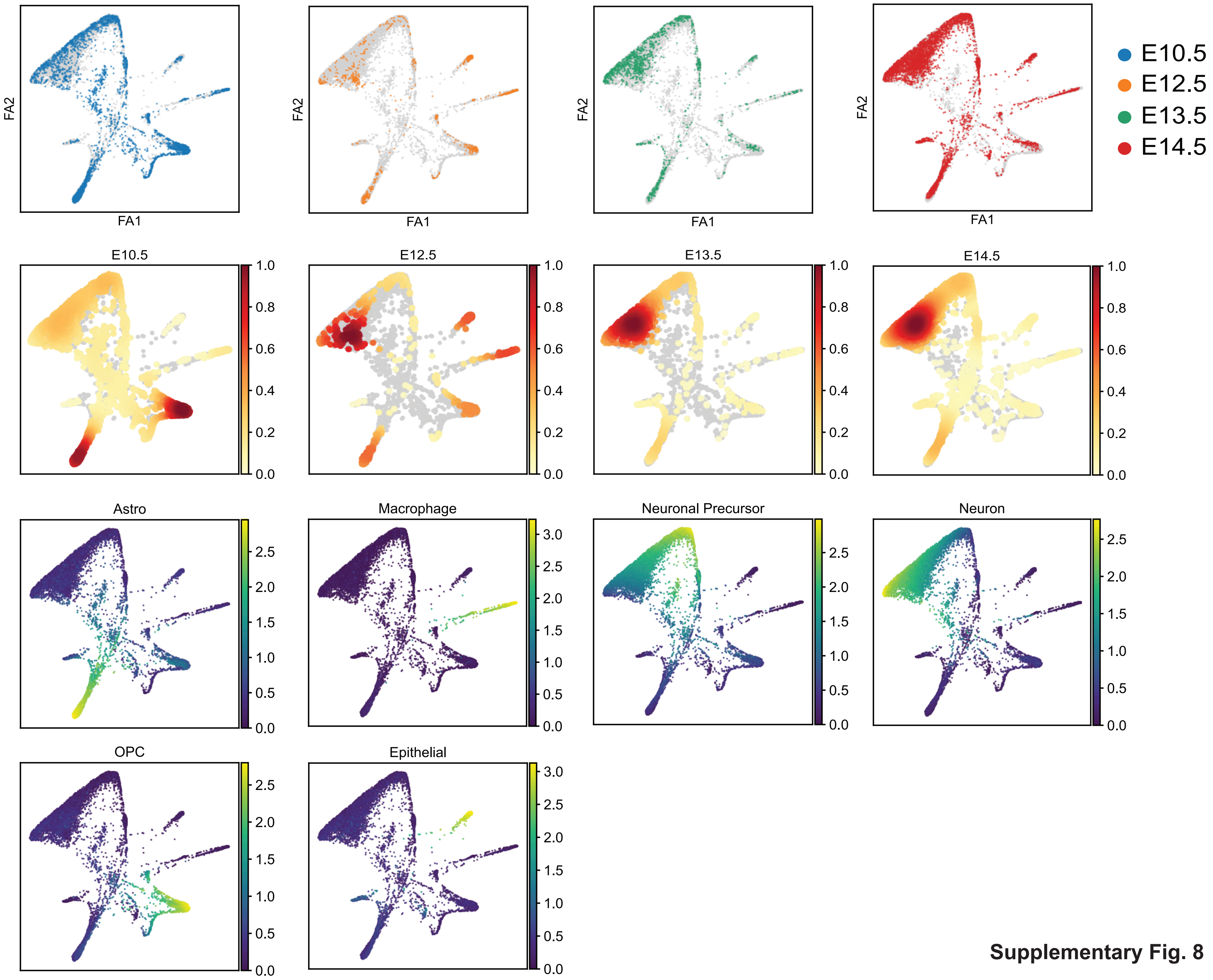


**Supplementary Fig. 8**

Supplementary Fig. 9

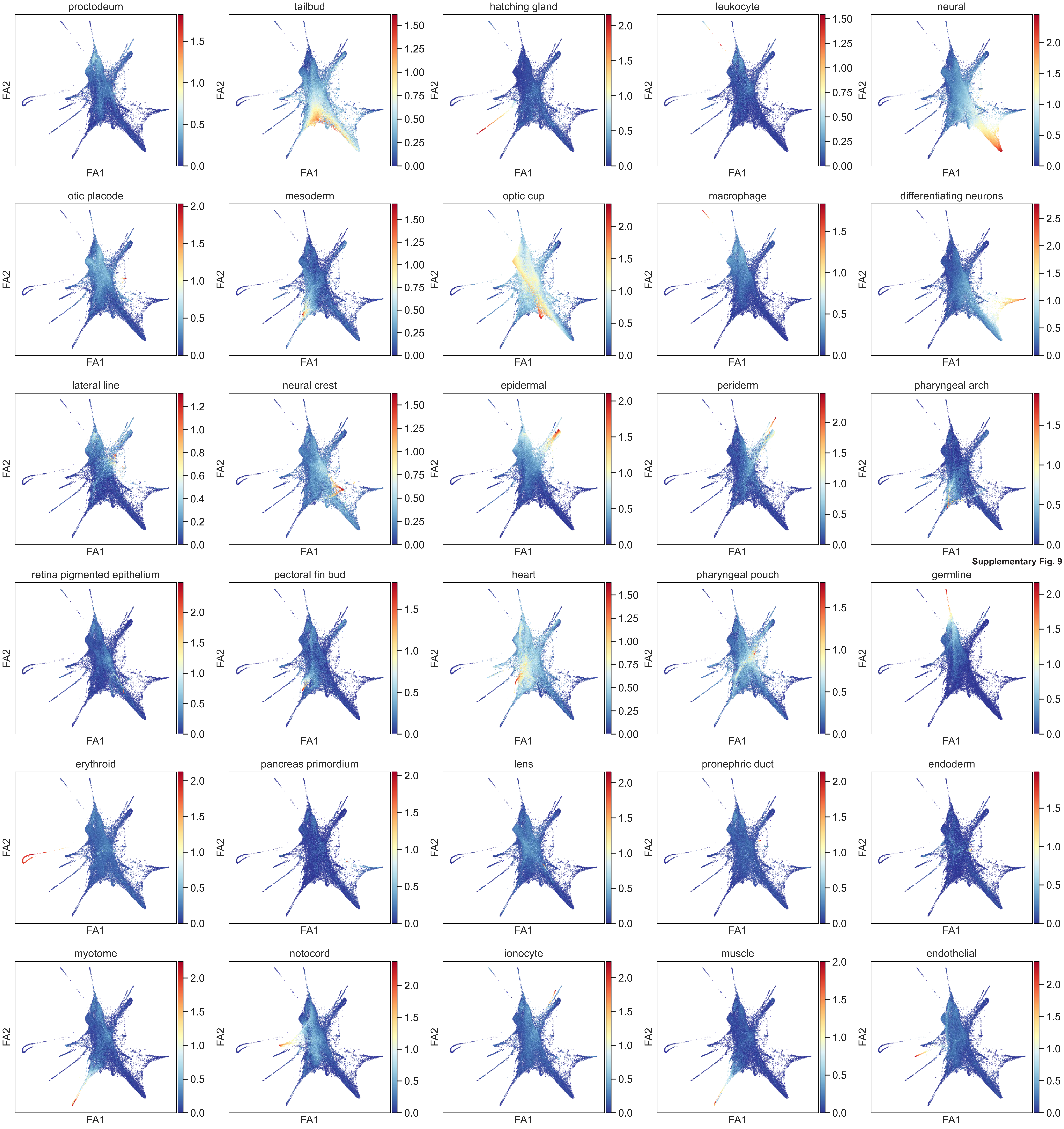

### Supplementary Fig. 10

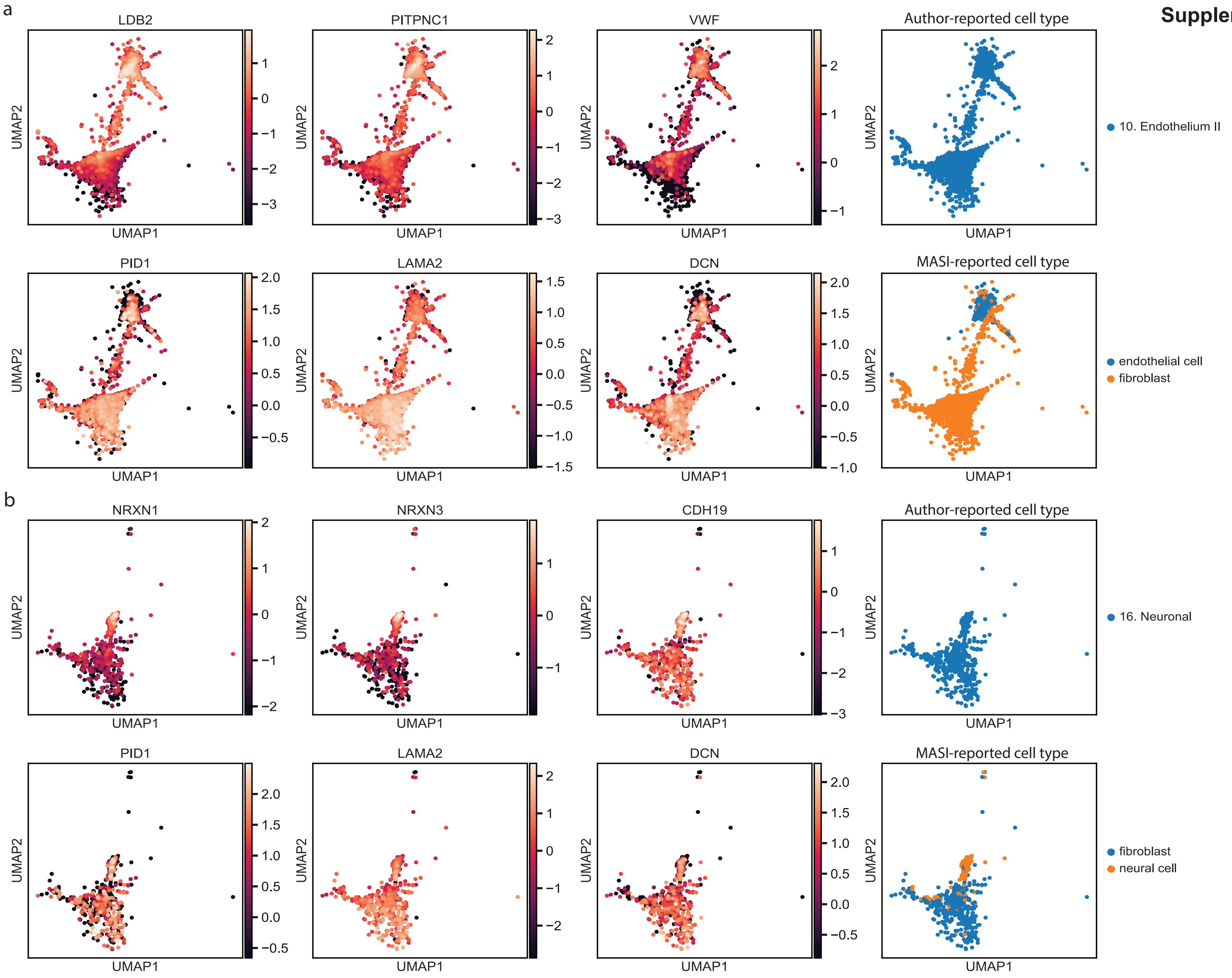

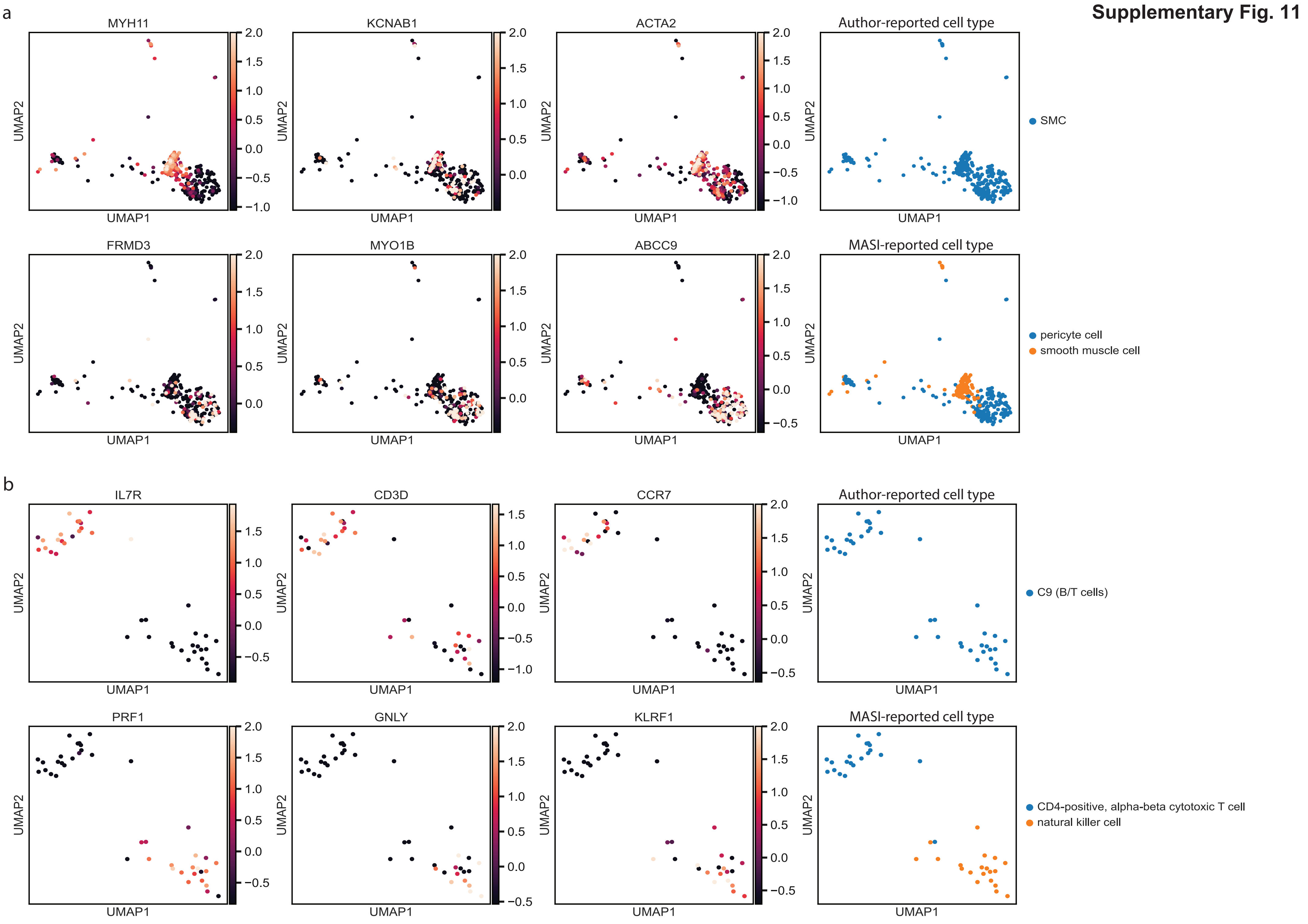
a
MYH11
KCNAB1
ACTA2
Author-reported cell type
SMC
FRMD3
MYO1B
ABCC9
MASI-reported cell type
pericyte cell
smooth muscle cell
b
IL7R
CD3D
CCR7
Author-reported cell type
C9 (B/T cells)
PRF1
GNLY
KLRF1
MASI-reported cell type
CD4-positive, alpha-beta cytotoxic T cell
natural killer cell
UMAP1
UMAP2

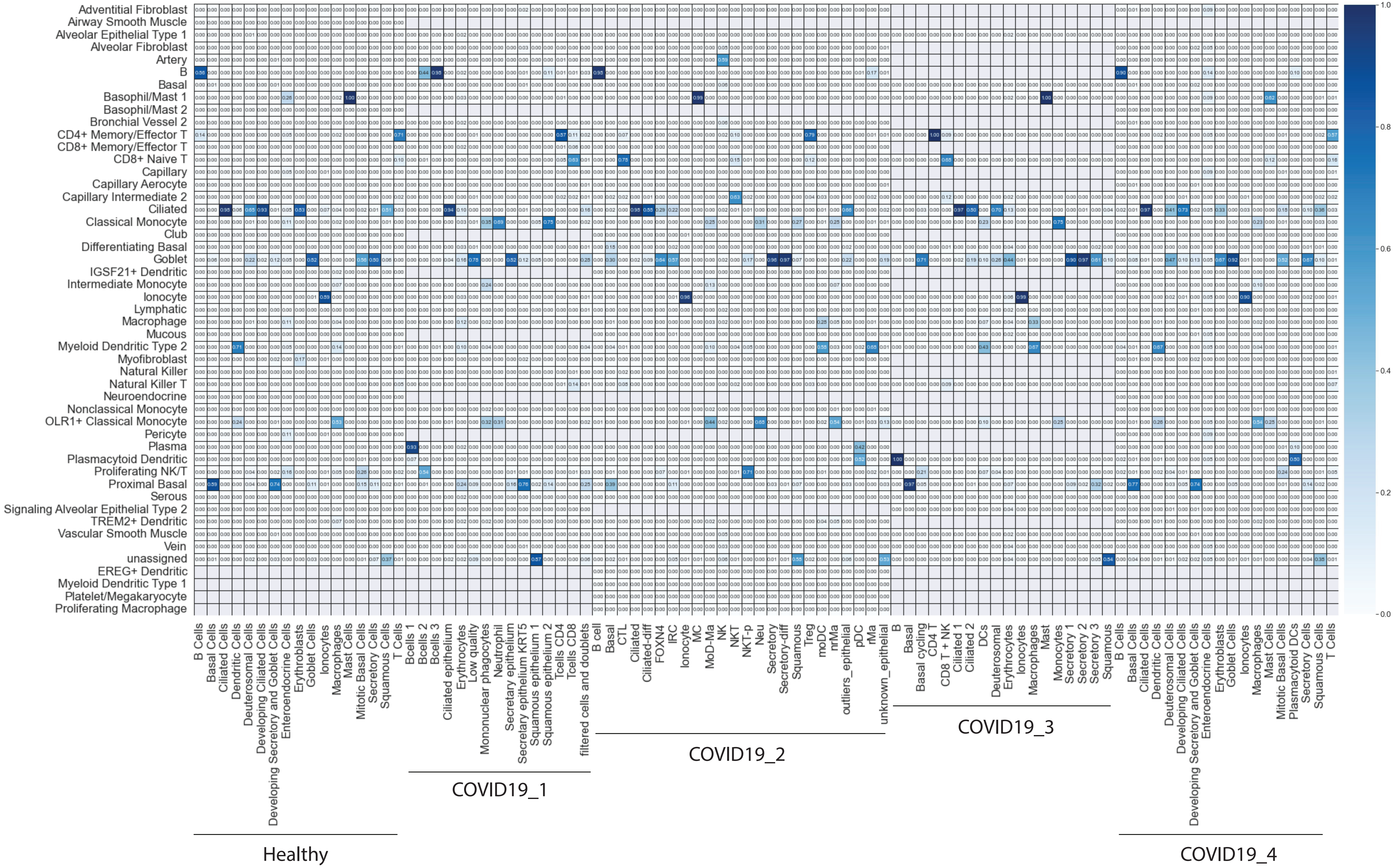


**Supplementary Fig. 12**

### Supplementary Fig. 13

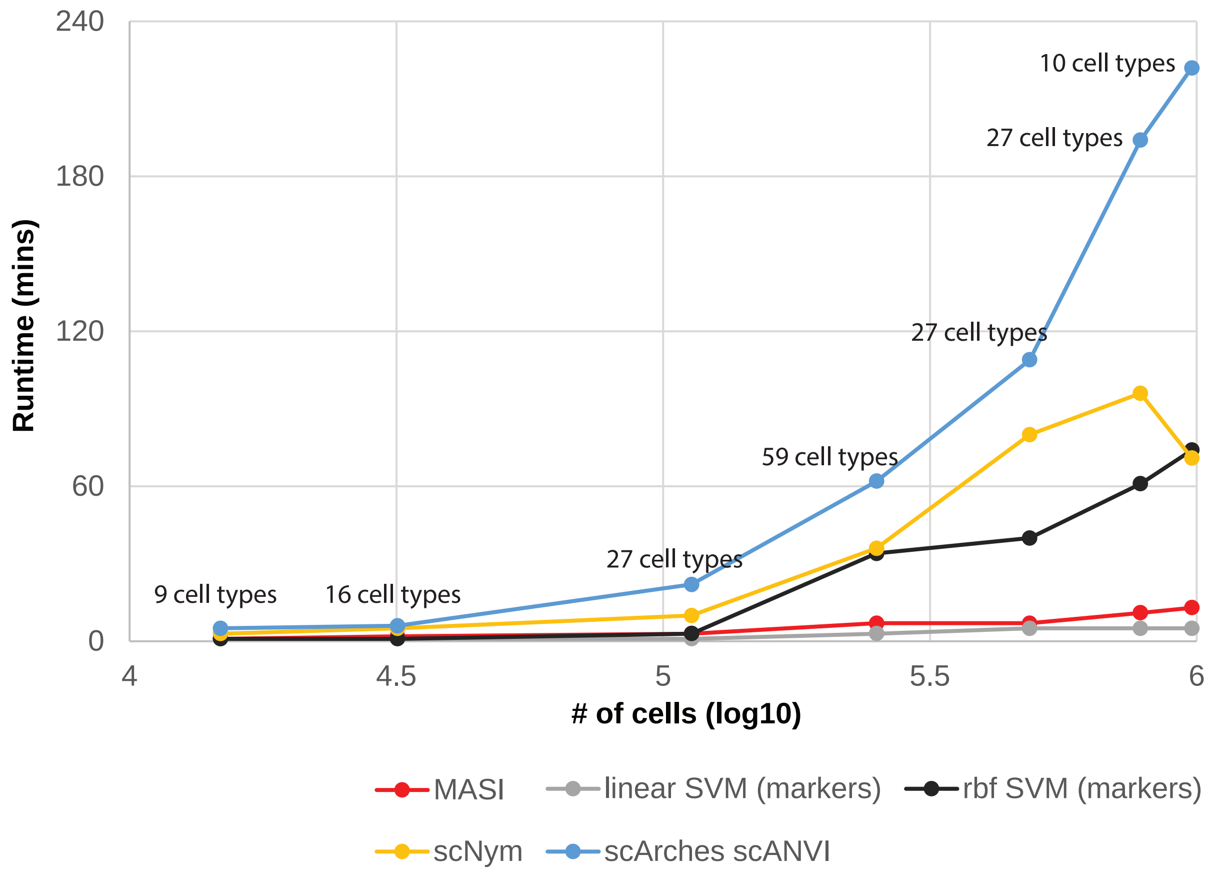

Supplementary Fig. 1: Illustration of 10 analysis pipelines for scRNA-seq data. Colored boxes highlight specific practices in a pipeline.

Supplementary Fig. 2: Benchmarking of impacts of 10 analysis pipelines on batch correction. a, UMAP visualization for cell-type preservation and batch mixing. Cells are colored according to author-reported cell-type annotation (left) and batch id (right). b, Evaluation of batch correction. Cell-type silhouette score (column) measures how well a pipeline perverse cell-type variation, and batch entropy mixing score (row) quantifies how well a pipeline mixed cells from different batches. Dots located in the top right should present good integration outcomes.

Supplementary Fig. 3: Benchmarking of impacts of 12 DE tests on MACA-based cell-type annotation. Cell-type markers are identified from a reference data using a specific DE test. Then, MACA annotates target data with markers identified by this specific DE test. Macro F1 score is reported to show how compatible the DE test is to MACA-based cell-type annotation.

Supplementary Fig. 4: Visualization of integration by scNym and scArches. Cells are colored according to method-reported cell-type annotation (left), author-reported cell-type annotation (middle), and batch id (right).

Supplementary Fig. 5: Hierarchical analysis of cell type. Spearman correlation is calculated with integrated representation by MASI, scNym, and scArches. Then, the dendrogram is constructed with “ward” as criterion.

Supplementary Fig. 6: Comparison of annotation resolution for MASI, SCCAF, and combination of SCCAF and MASI. 10X data is used as reference for label transferring. Cluster identification through SCCAF is based on 12-dimension cell-type score matrix. SCCAF is applied to MASI-reported annotation to further identify subtypes. ARI and NMI are calculated by comparing method-reported annotation with author-reported annotation.

Supplementary Fig. 7: Integration of scRNA-seq and Slide-seqV2 by MASI. scRNA-seq is used as reference and Slide-seqV2 data is annotated according to cell type identified in the reference. Markers for principal cells, endothelial tip and oligodendrocyte are selected for visualization, shown below cell-type annotation.

Supplementary Fig. 8: Integrative lineage analysis for multi-condition mouse embryo brain tracking study. Cell density, cell-type score, and batch id for mouse embryo brain samples under different conditions are visualized separately through the first two ForceAtlas2.

Supplementary Fig. 9: Visualization of 30 cell-type scores in developing zebrafish embryo. Cells from both Wagner et al and Farrell et al data are jointly shown in the first 2 ForceAtlas2.

Supplementary Fig. 10: Comparison of MASI-reported and author-reported annotations in Tucker et al data.

a, Comparison for author-reported endothelial cells. Expression of marker genes for endothelial cell (upper) and fibroblast (lower) are visualized in the first 2 UMAPs. b, Comparison for author-reported neural cells. Expression of marker genes for neural cell (upper) and fibroblast (lower) are visualized in the first 2 UMAPs.

Supplementary Fig. 11: Comparison of MASI-reported and author-reported annotations in Wang et al and Cui et al data. a, Comparison for author-reported smooth muscle cells in Wang et al data. Expression of marker genes for smooth muscle cell (upper) and pericyte (lower) are visualized in the first 2 UMAPs. b, Comparison for author-reported B/T cells in Cui et al data. Expression of marker genes for CD4+, alpha-beta cytotoxic T cell (upper) and natural killer cell (lower) are visualized in the first 2 UMAPs.

Supplementary Fig. 12: Confusion matrix of MASI-reported annotations against author-reported annotations in COVID19 datasets.

Supplementary Fig. 13: Runtime of 5 cell-type annotation methods. Datasets used in study are arranged from the least number of cells to the largest number of cells. The total number of unique cell types are marked on the top of runtime result.

Supplementary Table 1: Comparison of MASI running on two devices, workstation equipped with 10 cores Intel Xeon Silver 4210 and 64GB memory (high-end device), and personal laptop equipped with 4 cores Intel i7-8550U and 16GB memory (basic device).

Supplementary Table 2: Description of data used. Brief description, including species, number of cells and cell types, GEO accessions, etc. are listed.
